# The tuxedo sea urchin *Mespilia globulus*: A fast-developing and tractable model for genomic and developmental biology

**DOI:** 10.64898/2026.07.23.740292

**Authors:** Omar Matar, Marie Emilie Maeland, Lucas King, Elise Parey, Grace Birkett, Clara Santangelo, Laura Piovani, Jamie Craggs, Jeffrey R. Thompson, Nathalie Oulhen, Gary Wessel, Ferdinand Marlétaz

## Abstract

Sea urchins are pivotal models in cellular and developmental biology, but their biphasic life cycle with an extended larval life limits the ability to study post-metamorphosis and adult characters. Here, we introduce a genomically-enabled model system, the tuxedo urchin *Mespilia globulus*, which has rapid access to late life stages in sea urchins. We describe how we cultured *M. globulus* in a landlocked marine facility, raised larvae under artificial conditions and closed their life cycle. We established the experimental tractability of *M. globulus*: we labelled transcripts by hybridization chain reaction (HCR), and knocked out pigmentation genes to produce albino larvae using CRISPR/Cas9. We generated chromosome-scale genome assemblies for two individuals representing both sexes and two color morphs (red and blue), and compared the organisation of the 21 chromosomes of *M. globulus* with that of other camarodont echinoid models. We determined that *M. globulus* showed a conservative gene repertoire lacking the gene family expansions seen in other camarodont sea urchins. We annotated the complement of genes associated with pigmentation, immune and nervous systems and profiled their expressions in tissues and organs. Finally, we surveyed sex-related regions in genomes using genome assemblies and resequencing data, finding no evidence of heteromorphic sex chromosomes in *M. globulus*. Our findings highlight the accessibility of this new sea urchin model for studying the metamorphosis and adult biology of sea urchins.

## Introduction

Sea urchins are pivotal models for embryology as they spawn millions of transparent eggs, enabling embryologists to access and manipulate developmental processes (Hörstadius, 1973; McClay, 2011). Many key discoveries have been made using sea urchins as experimental models, including mechanisms of fertilization and morphogenesis, cell cycle control, and gene regulatory processes leading to cell type specific differentiation (Ernst, 2011; Hörstadius, 1939; McClay, 2011). Sea urchin eggs and embryos can be microinjected and experimentally manipulated, which has been instrumental in applying gene interference techniques to reconstruct the gene regulatory networks controlling animal development (Davidson, 2006; Davidson and Britten, 1973; Franks et al., 1988; Heyland et al., 2014; Oliveri et al., 2008). Sea urchins transition from bilateral embryos and larvae to a pentaradial adult body plan during metamorphosis (Arenas-Mena et al., 1998; Hyman, 1955). In the popular sea urchin models such as *Strongylocentrotus purpuratus* or *Hemicentrotus pulcherrimus*, the long larval and juvenile stages (>18 months) complicate investigation of the process underlying this transition (Smith et al., 2008; Yajima, 2007). Similarly, echinoderms possess many unique organismal traits, such as the water vascular system for function of the tubefeet, as well as diversity in spine morphology and pigmentation patterns, for which developmental and genetic underpinnings remain to be understood (Arnone et al., 2015; Wise et al., 2024). For instance, previous studies have suggested that the same genes could be involved in the establishment of both bilateral and pentaradial body plans, first during gastrulation and then during larval metamorphosis (Gildor and Ben-Tabou de-Leon, 2015; Li et al., 2020). The long generation time in *S. purpuratus, H. pulcherrimus* though, has severely limited the development of genetic resources, such as transgenic lines.

Since the original release of the *S. purpuratus* genome (Sea Urchin Genome Sequencing Consortium et al., 2006), multiple sea urchin genomes have been sequenced to chromosome-scale level, including *Lytechinus variegatus* (Davidson et al., 2020), *Lytechinus pictus* (Warner et al., 2021), *Hemicentrotus pulcherrimus* (Komoto et al., 2024) and *Paracentrotus lividus* (Marlétaz et al., 2023), which all belong to the camarodont clade, a highly diverse order of regular euechinoids (Mongiardino Koch et al., 2022). Comparative analysis of these genomes has revealed that the ancestral bilaterian chromosomes are globally maintained in the ∼21 chromosomes of sea urchins, but conversely that the intrachromosomal local gene order evolved quickly, particularly compared to vertebrates (Marlétaz et al., 2023; Parey et al., 2024). While some cytogenetic studies suggest heteromorphic sex chromosomes in *P. lividus* (Lipani et al., 1996), and while sex determination has been described as polygenic in sea cucumber (Jiang et al., 2024), the genomic determinant of sex differentiation remains undeciphered in sea urchins.

The success of Cas9 gene knock-ins, Cas9 gene knock-outs and Minos-mediated gene integrations expands the scope of experimentation possible in sea urchins (Caccavale et al., 2026; Lin et al., 2019; Oulhen et al., 2023; Wessel et al., 2021). Researchers are now capable of testing gene function in transgenerational studies, and making various genetic strains using fluorescent reporters for mechanistic studies of the genes involved in, for instance, fertilization and pigmentation (Jackson et al., 2024; Lin et al., 2019; Oulhen et al., 2023; Wessel et al., 2020). A limitation of these systems, however, is the husbandry attention required to raise the animals to gravid adults, as well as the time and effort it takes over protracted intervals of time, with up to 18 months to reach gravid adults for transgeneration studies (*S. purpuratus* or *H. pulcherrimus*). Recently, *Lytechinus pictus,* a species from the eastern Pacific coast, has been introduced as a genetically tractable sea urchin thanks to its faster post-embryonic development and earlier sexual maturity (Nesbit et al., 2019; Nesbit and Hamdoun, 2020), which has permitted, in particular, the establishment of transposon-based transgenic lines (Jackson et al., 2024). Cryopreservation protocols for sperm and various stages of embryogenesis make it possible to share animal strains between labs expanding the genetic capabilities of sea urchins (Vacquier and Hamdoun, 2025, 2024). Together with genome sequencing, these recent advances have the potential to enable the development of high-throughput genetics approaches in sea urchin models (Lee et al., 2025). However, the majority of transgenerational experimentation has transpired in marine labs, with ready access to natural sea water from whence the animal came. In cases where access to natural sea water is impossible, a rapidly developing sea urchin with robust growth and maturity characteristics in closed aquarium systems is a necessary addition to the repertoire of model systems for unconfined experimentation.

With these goals in mind, we introduce *Mespilia globulus*, a sea urchin species native to the tropical waters of the Indo-Pacific region, as an additional model system with a rapid developmental program and attractive properties to study larval development and adult characteristics of echinoid sea urchins. *M. globulus* is a popular species in the aquarium industry and has excellent properties in tropical aquariums where it has been bred for many years in farming operations that reduce genomic heterogeneity and enhance sustainability of the species and the ecosystem. Its co-culturing in marine systems can improve coral maintenance and, in doing so, it contributes to a healthy marine aquarium environment (Craggs et al., 2019).*M. globulus* belongs to the infraorder Temnopleuridea, and therefore represents an earlier diverging lineage within the camarodonts than most other model sea urchins (**Figure 1B**) (Mongiardino Koch et al., 2026). We report how this species can be cultivated and its life cycle enclosed within a land-locked, minimal marine facility, and describe how we validated an array of experimental techniques, such as microinjection, HCR-FISH staining and CRISPR/Cas9 gene knock-outs. Then, we report multiple genome sequences of *M. globulus* using long-read technology and proximity-ligation based sequencing from distinct color morphs, including both male and female sexes, providing insights into sex determination mechanisms in sea urchins.

**Figure 1.**
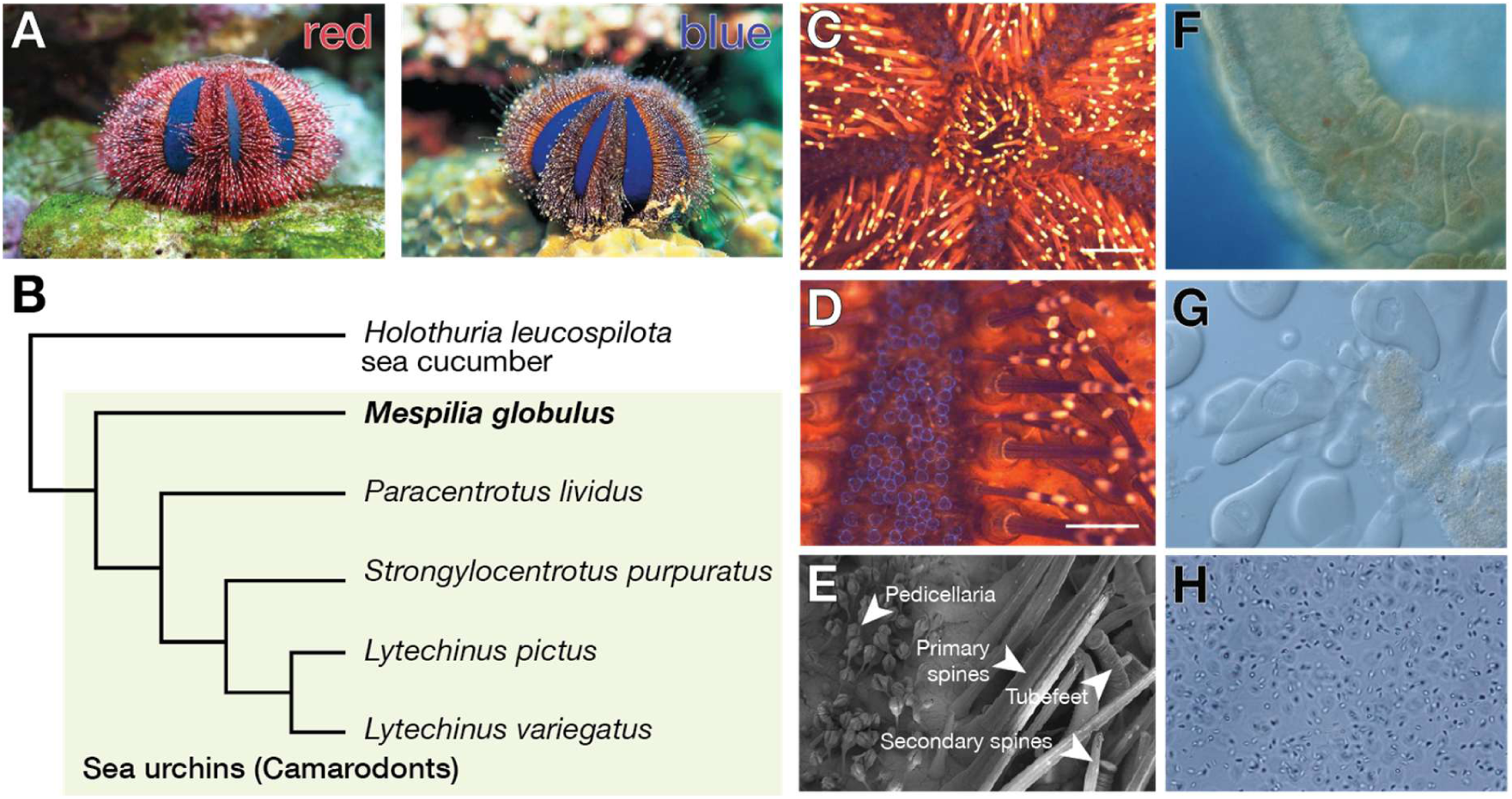
The pigmented and reproductive structures of *Mespilia globulus.* (**A**) Red and Blue distinct color morphs. (**B**) Phylogenetic position of *M. globulus* among echinoid sea urchins (Mongiardino Koch et al., 2026). (**C**) Banded spines and blue ensheathed pedicellaria visible in the aboral region using light microscopy (**C** and **D**) and SEM (**E**) (scale bar 500µm). (**D**) The ovaries are arranged in tubular structures with closely opposed oocytes. (**F** and **G**) The developing oocytes are of very similar sizes, here misshapen from the ovary dissection. (**H**) Sperm have the characteristic triangular shaped head and long flagellar tail.

## Results

### *Mespilia globulus,* a model for pigmentation, reproductive and developmental biology

*M. globulus* is striking in appearance: red and blue color morphs make the species an attractive model to study color patterns including striped spines, clear delineations of ambulacral and interambulacral regions and deep blue pedicellaria (**Figure 1A**). *M. globulus* is gonochoristic with a distinct ovary and testes (**Supp. Figure 1**). The ovary has the characteristic lobulated feature of sea urchins with abundant oocytes containing a large germinal vesicle with a prominent nucleolus, and a pronucleus following meiosis. The ovaries appear to have synchronously developing oocytes as seen in some other species of echinoids (e.g. *Diadema antillarum*)(Iliffe and Pearse, 1982). The sperm are also of classic morphology with a triangular shaped head and long flagellar tail (**Figure 1H**). These particular characteristics also make *M. globulus* an attractive system to study sea urchin reproductive biology.

We developed a reproducible and scalable culture system to maintain *M. globulus* where adult urchins were housed in custom-designed artificial seawater (ASW) tanks, featuring continuous flow-through filtration, temperature regulation, light cycling to simulate tropical marine conditions and parameter monitoring (temperature, pH) (**Supp. Figure 2**).

In this system, we investigated the pace and robustness of *M. globulus* embryonic and post-embryonic development until settlement (**Figure 2A**). After spawning and fertilization, embryos were transferred to large culture vessels fitted with paddle stirrers and gentle aeration systems to prevent sedimentation and to maintain water circulation (**Supp. Figure 2C**). We found that the development of *M. globulus* is rapid, reaching metamorphosis in around 14 days post-fertilization (dpf) (**Table 1**). This is markedly faster than the 30-45 dpf typically required for metamorphosis in traditional temperate urchin models such as *P. lividus, H. pulcherrimus,* and *S. purpuratus,* and comparable to reports of *Lytechinus pictus* (Nesbit et al., 2019) (**Table 1**). At the onset of metamorphic competency, larvae underwent spontaneous settlement within the same beakers, and post-settlement juveniles were carefully transferred to rearing boxes (**Supp. Figure 2B**). We raised juveniles to sexual maturity (see **Methods**) in 5 months, effectively closing the life cycle, which highlights the model system’s potential of this species, especially in a landlocked research laboratory.

**Figure 2.**
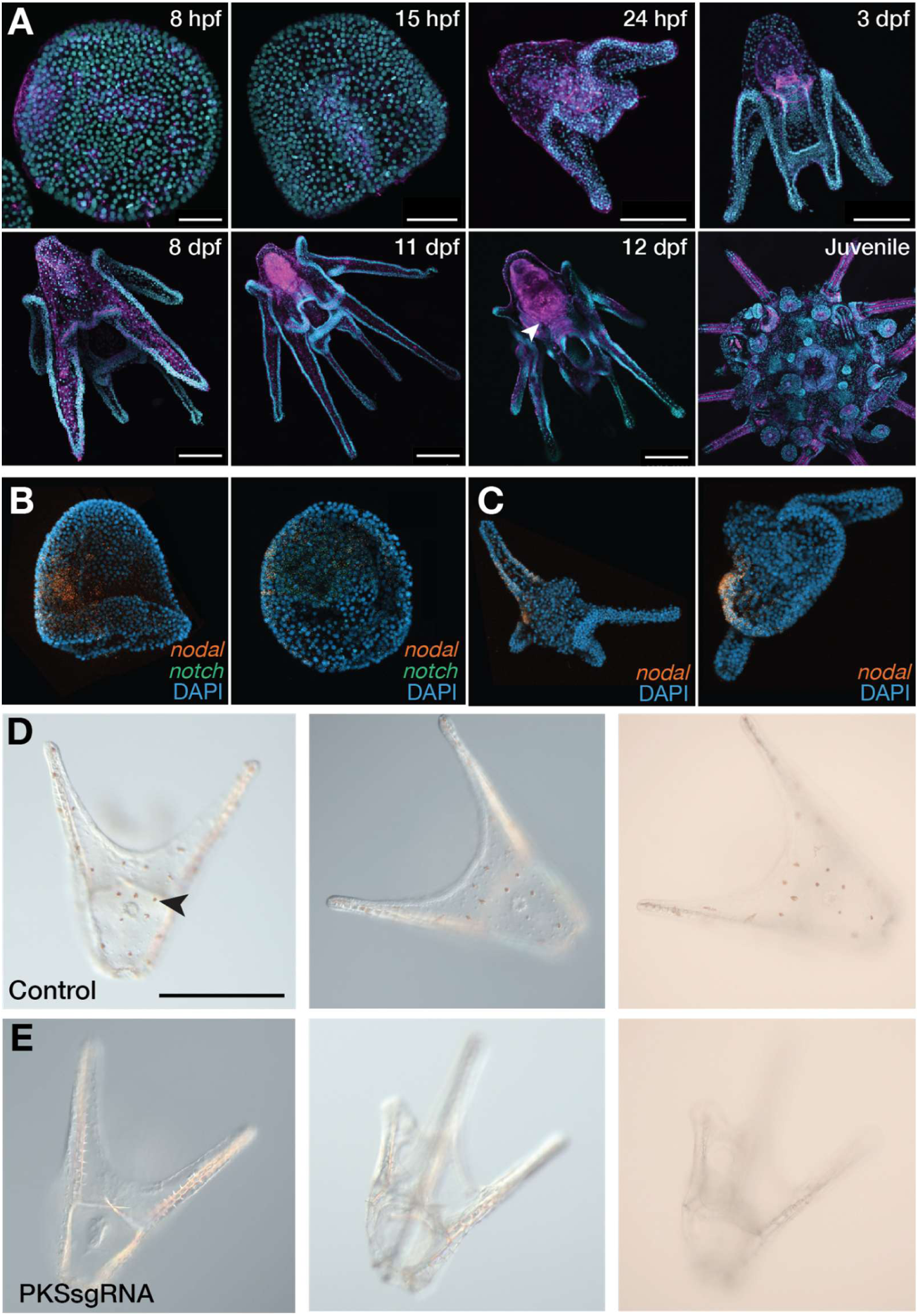
Development, HCR staining and CRISPR/Cas9 in *M. globulus*. (**A**) Confocal images of the development of *M. globulus* from early gastrulation through to an early juvenile. Nuclei in cyan and Muscle fibers in Magenta. (**B**) HCR stainings in early gastrula of the *nodal* and *notch* transcripts. (**C**) HCR stainings of *nodal* in pluteus larva. In (**B,C),** left and right panels correspond to lateral and apical views and nuclei are counterstained with DAPI. (**D**) control of CRISPR/Cas9 knockout showing the pigment cells in an early pluteus (arrowhead) in three different imaging modalities. (**E**) Cas9 inactivation of the pigmentation gene polyketide synthase (PKS) resulting in albinism, a phenotype with a lack of pigmentation as seen in the same imaging modalities.

**Table 1:**
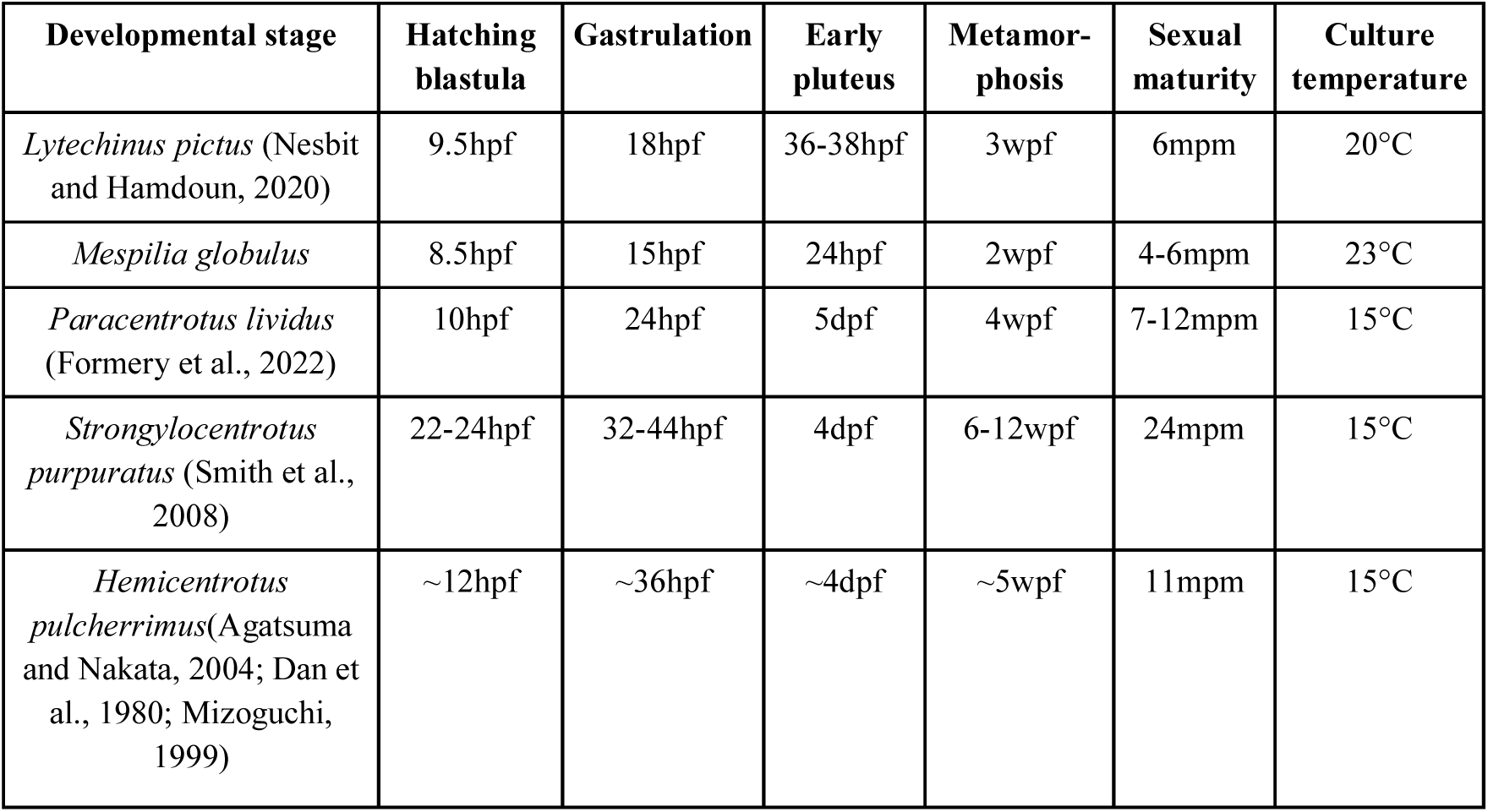
Developmental rate of different urchin species. hpf: hour post fertilization, dpf: day fertilization, dpf: week fertilization, mpm: month post metamorphosis. Sexual maturity corresponds to the ability to make gametes.

### Experimental tractability of *M. globulus*: HCR labeling and CRISPR/Cas9

We further characterised the anatomy of embryonic, post-embryonic and juvenile *M. globulus* using a nuclear stain (DAPI) and an actin stain (Phalloidin). These labels reflect the clarity in developmental features, including the growing adult rudiment (**Figure 2A**, 12dpf, arrowhead). While previous attempts to culture tropical urchin species have faced challenges particularly in larval survival, achieving settlement, and maturation with *M. globulus* outside of natural seawater environments represent a step forward in echinoderm system accessibility (Craggs et al., 2019).

To assess the use of *M. globulus* as a tractable tropical model system, we tested and adapted several essential protocols to survey gene expression and function during development. We first attempted to apply *in situ* hybridization chain reaction (HCR) which is a powerful approach for multiplexed *in situ* gene expression monitoring, successfully applied in diverse animal models, including sea urchin (Choi et al., 2016). We profiled two key developmental regulators, *Nodal* and *Notch*, during embryogenesis (**Figure 2B,C**). Nodal expression was observed in the presumptive oral ectoderm, consistent with its conserved role in establishing the left-right (L/R) axis and oral-aboral polarity. Notch was detected in regions consistent with signaling centers involved in mesodermal specification and later-stage rudiment patterning (Duboc et al., 2005). These expression domains mirror those described in well-characterized echinoid systems, supporting a conservation of molecular developmental program in *M. globulus*, and particularly the role of *Nodal* in the left-sided restriction of rudiment formation (Duboc et al., 2005).

Next, we tested the facility of Cas9 function in *M. globulus* by injecting eggs with Cas9 protein containing sgRNAs targeting the large iterative enzyme polyketide synthase (PKS) (**Figure 2D** and **E**). PKS synthesizes polyketides, some of which are responsible for making echinochrome and other spinochrome pigments within sea urchins (Calestani et al., 2003). Previous reports documented that larvae lacking PKS function are albinos (Oulhen and Wessel, 2016), as are adults (Wessel et al., 2020), and the larvae tested here with Cas9/sgRNA developed perfectly well, but were albino (**Figure 2E**).

### Genomes of red and blue morphotypes

We sequenced the genomes of two morphotypes and both sexes of *M. globulus*: a red female and a blue male sea urchin. These color morphs differ in the amount of red pigment present in the banded spines and test of the adult (**Figure 1A**). We used PacBio HiFi long-read sequencing in both cases and proximity-ligation based scaffolding yielded chromosome-scale assemblies (**Figure 3A,B** and **Supp. Figure 3**).

**Figure 3.**
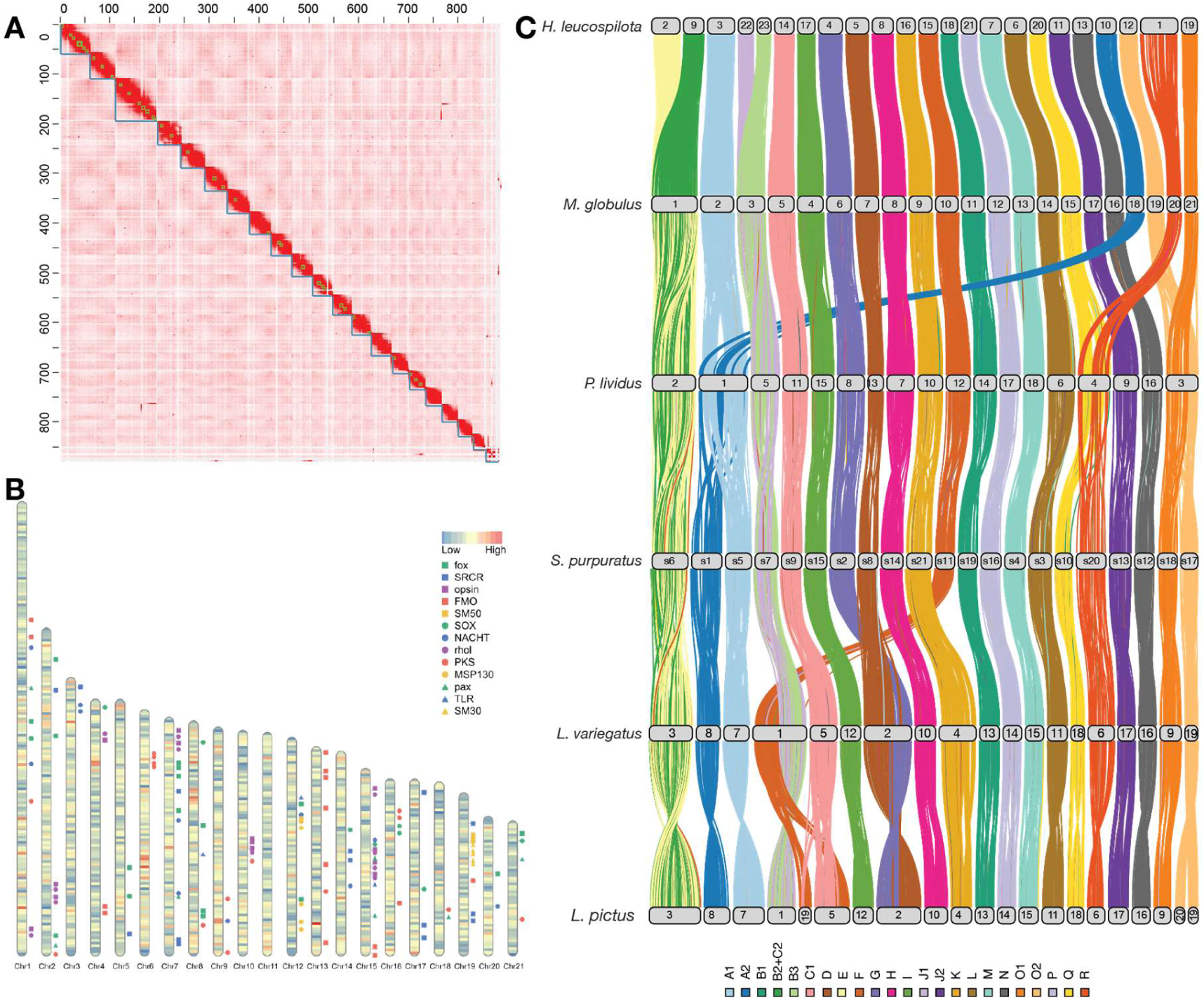
The genome of *Mespilia globulus* and the evolution of sea urchin genomes. **(A**) HiC contact map highlighting the 21 assembled chromosomal scale scaffolds. (**B**) Gene density in the chromosomes of M. globulus indicated several gene duplication hotspots associated with gene families involved in organismal novelties of sea urchins (see Figure 5). (**C**) Evolution of sea urchin genome architecture using single copy orthologues annotated for ancestral echinoderm linkage groups.

The blue male and red female have a total assembly size of 977Mb and 870Mb with a N50 of 41.8Mb and 39.8Mb, respectively. Both assemblies have high completeness scores with >99% of complete BUSCOs. Both assemblies have 21 main chromosome-sized scaffolds with strong internal HiC contacts, which is comparable to chromosome number in other sea urchin species (**Figure 3A**). The integrity of these chromosomal sets was also further tested by analysing their synteny (see below). We numbered the chromosomes based on their size in the blue male assembly, and matched them to the red female assembly for convenience in comparisons using whole genome alignment (**Supp. Figure 4**). We impute the marginal differences in the size of chromosomes across assemblies to differences in coverage and polymorphism between the two sampled populations (see below). Genome annotation, performed using existing and newly generated transcriptomes (**Supp. Table 2**), identified 21,039 protein-coding genes, of which 75% included PFAM domains.

We compared the genomes of the two color morphs using whole-genome alignments (WGA) and orthology-based Oxford dotplots (**Supp. Figure 4**). We found that the two assemblies are structurally identical with one-to-one correspondence between chromosomes across morphotypes, and that gene order is essentially colinear, with only a few signs of rearrangement within chromosomes. For instance, chromosomes 1, 7 and 15 show limited inversions in terminal regions (**Supp. Figure 4**). As genomes are globally colinear, we estimated nucleotide divergence between the two morphotypes and found this to be 3.1% from our alignments. In comparison, within-population polymorphism levels were 1.3% for both the red morphotype and the blue morphotype. The phenotypic difference between these morphotypes does not appear associated with extensive genomic difference, both at the structural and nucleotide level, and therefore we investigated below whether more localised variation in regulatory or coding-sequences (*e.g.* PKS gene) could be responsible for color differences.

### The genomics of sex determination

We next asked whether any of the observed differences between the genomes could be related to the presence of heteromorphic regions involved in sex determination. Earlier reports described a heteromorphic sex chromosome in *P. lividus*, another camarodont species (**Figure 1A**) with marked difference in size and morphology for one of the biggest pairs of chromosomes (Lipani et al., 1996). We did not observe marked size differences between male and female for any of the large chromosome pairs (**Supp. Figure 5A** and **Supp. Table 3**). The two smallest chromosomes showed higher difference in size (18% and 24% for chr. 20 and 21, respectively) but the difference mostly corresponds to low complexity and gene poor terminal regions **(Supp. Figure 5D**).

As such differences can be explained by differences between individuals or populations, we generated 30× short read coverage for a distinct pair of male and female individuals from the blue morphotype to investigate possible coverage discrepancies (**Figure 4**). We calculated a genome-wide normalised sex coverage ratio (M/F) on both the male and female assemblies. None of the chromosomes (**Figure 4A**) and few broad chromosomal segments (**Figure 4B**) showed a skewed ratio consistent with a sex-specific identity (plus/minus 0.5 ratio). Similarly, the regions specific to the female assembly also appear to not show a consistent ratio compatible with a sex-bias. The most intriguing locus is a broad region of ∼20Mb on chr. 4, possibly corresponding to a full chromosome arm, that shows a limited but consistent skew in female coverage (av. −0.15) on both male and female genomes assemblies (**Figure 4C**). We therefore searched this region for any known genes associated with sex determination but did not find any classically conserved genes (*e.g. Dmrt*, *Sox*, *Sry*, Aromatase, *etc…*) (Capel, 2017) and similarly, we did not observe any sex-related GO term enrichment (**Supp. Figure 5** and **Supp. Table 4**).

**Figure 4.**
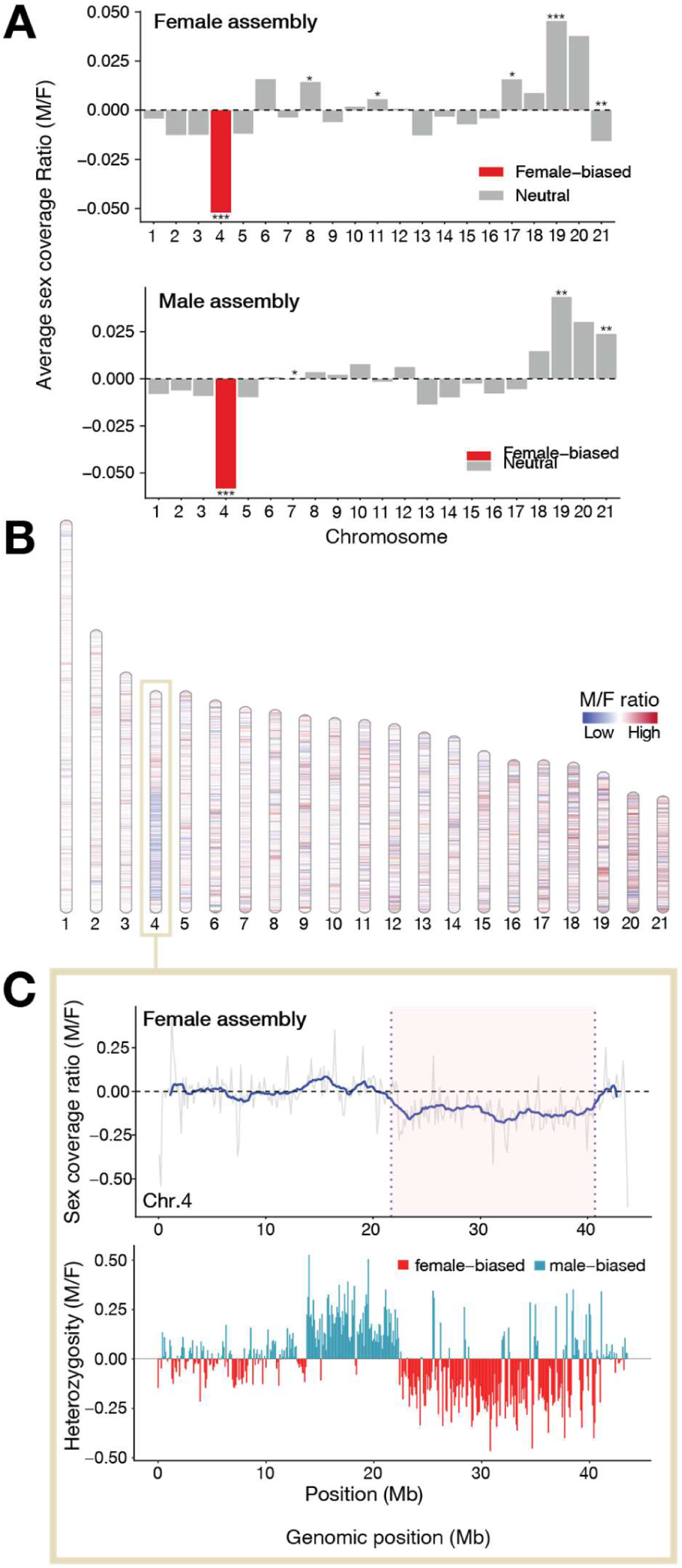
Genome-wide distribution of sex-biased coverage across chromosomes. (**A**) Average normalised sex coverage ratio (M/F) for each chromosome of the male and female assemblies. The difference in coverage was assessed using a Wilcoxon rank test with a FDR correction (*: p-value=0, ** P-value < 1e-5, * p-value <0.05) (**B**) Distribution of M/F sex ratio in the 21 chromosomes across genomic windows. Red indicates relative female enrichment, blue indicates relative male enrichment, and white indicates balanced coverage. (**C**) Region on a putative arm of chromosome 4 showing a female bias.

Overall, our observations point toward the absence of large-scale karyotypic differences between sex, and instead suggest a polygenic sex determination system, involving sex-related allelic variation. These allelic differences could be concentrated in specific regions, such as one arm of chr. 4, and explain the slight but consistent difference in mappability without full-scale sex ratio variation. Similar sex determination mechanisms have been reported in sea cucumbers, which are the sister-group to sea urchins (Jiang et al., 2024). Our findings do not directly invalidate the karyotypic observations made by Lipani *et al*. in *P. lividus* and it is possible that over the course of sea urchin evolution, this polygenic mechanism transitioned toward an heteromorphic system, suggesting that chr. 4 could constitute a proto-sex chromosome (Graves, 2015).

### Evolution of genomic architecture across sea urchin species

We examined the chromosomal correspondence across five sea urchin species and a sea cucumber (**Figure 3C, Supp. Figure 6** and **Supp. Table 5**). Sea cucumbers are the sister-group to echinoids in the echinoderm tree (Telford et al., 2014) and their 23 chromosomes match exactly the ancestral echinoderm linkage groups, which only diverge from the ancestral bilaterian units by the fusion- and-mixing B2⊗C2 (Parey et al., 2024) (using the algebraic notations of (Simakov et al., 2022)). We traced the evolution of these ancestral units in sea urchin chromosomes, and identified two fusion-and-mixing events shared by all euechinoids examined: E⊗(B2⊗C2) and J1⊗B3. The genome of the tuxedo urchin does not exhibit any further rearrangements in addition to those. This is also the case of *S. purpuratus*, and is therefore in one-to-one correspondence with *M. globulus*. Conversely, *P. lividus* shows 3 more fusions, some of them fairly recent and not yet followed by mixing (A1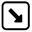A2; O1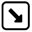O2; Q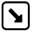R), as previously noted (Marlétaz et al., 2023). *Lytechinus* species share one recent fusion (D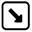G) and each shows a species specific fusion: (D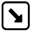(J1⊗B3)) in *L. variegatus* and (C1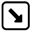F) in *L. pictus*. Notably, the genome assembled for *L. pictus* is based only on HiFi sequencing without Hi-C scaffolding and it is possible that some scaffolds indeed correspond to chromosomes (*e.g.* scaffold 19 and 20 in **Figure 3C**). These observations provide a valuable framework to compare different sea urchin models. They also highlight the relatively dynamic genomic evolution of echinoids, even if they present less rearrangements than the brittle stars, another group of echinoderms (Parey et al., 2024).

### Gene family evolution and sea urchins traits

Sea urchins are remarkable for their unique characteristics that play a key role in their adaptation to the environment, such as immune function, sensory biology, pigmentation and skeletogenesis (Smith et al., 2006; Wise et al., 2024). To determine whether changes in gene complement are associated with the evolution of these traits, we first examined a set of candidate genes involved in such processes and focused on possible changes in copy number (**Figure 3B**, **5A** and **Supp. Table 6**). Following this, we then performed a comprehensive reconstruction of changes in gene complement across sea urchins (**Figure 5B** and **Supp. Table 5**).

**Figure 5.**
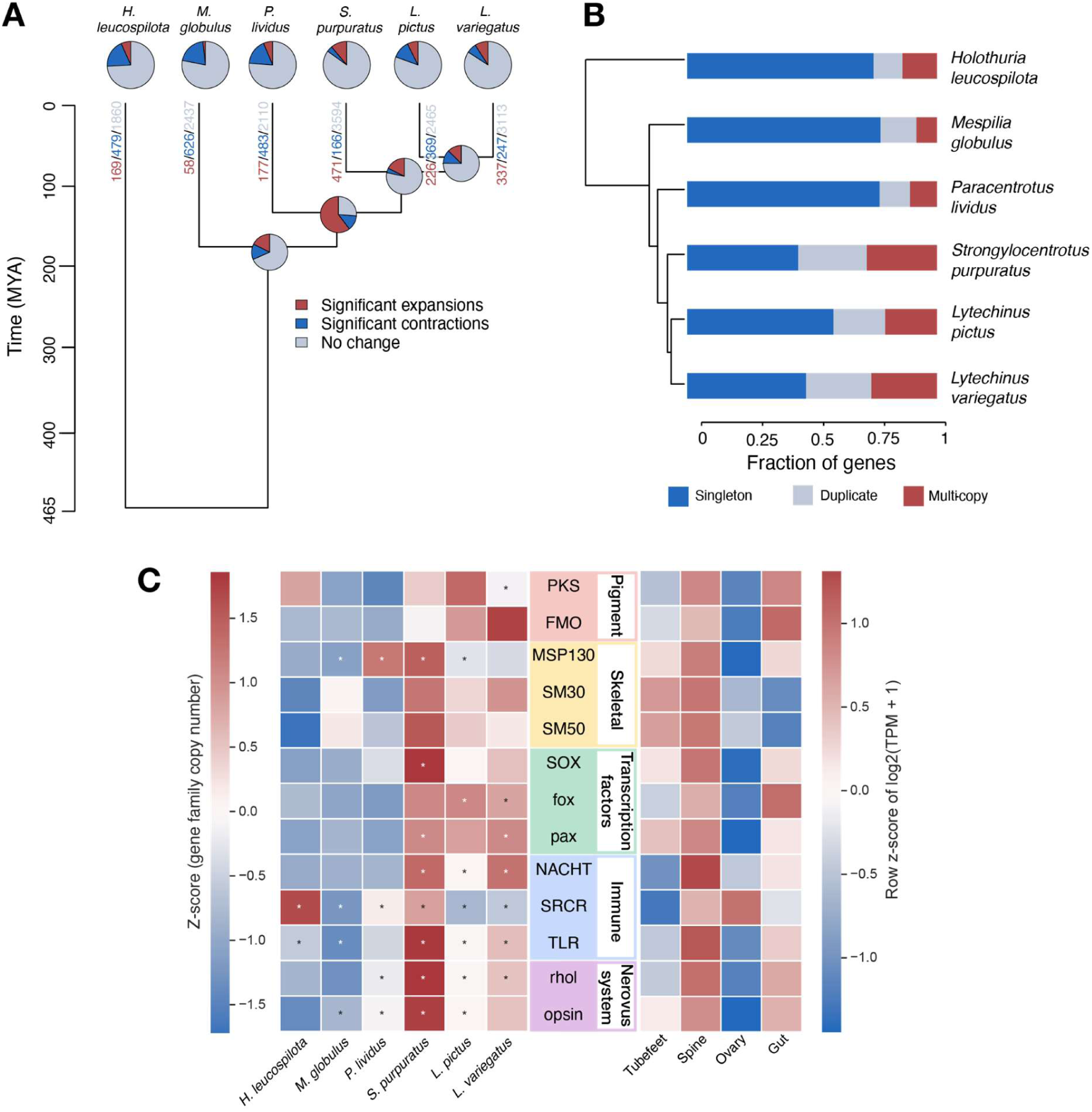
Gene family evolution and tissue expression of selected multifunctional gene families. (**A**) Ultrametric tree used for the CAFE analysis, with pie charts summarising expanded, contracted, and unchanged gene families at terminal and internal nodes. (**B**) Proteome composition, showing the proportions of singleton, duplicate, and multi-copy genes in each species. (**C**) Left, normalised heat map showing significant expansions and contractions (*) for each gene family from the cafe output; right, tissue expression of the same gene families across *Mespilia globulus* tissue transcriptomes.

As major differences in genome architecture are not associated with color morphs, we paid particular attention to a set of genes involved in pigmentation, including polyketide synthases (PKS) and flavin-containing monooxygenases (FMO), which are key enzymes in euechinoid biosynthesis (Wessel et al., 2020). We found that the FMO gene family showed variation in copy number among euechinoids, with pronounced expansions in *L. pictus* and *L. variegatus*, but not in *M. globulus* (**Figure 5C**), suggesting lineage-specific diversification of pigmentation pathways rather than a universal duplication trend across echinoids. Comparative analyses including earlier-diverging echinoids will be important for resolving the ancestral state of these pigmentation gene families.

Across specific functional categories (pigment, skeletal, transcription factors, immune, and nervous system; **Figure 5C**), comparisons of normalised gene copy numbers revealed strong phylogenetic biases in gene family dynamics. The majority of expansions occurred within the Odontophora (*S. purpuratus, L. pictus,* and *L. variegatus*), which together had broad gene family diversification across all of the targeted biological processes. Transcription factors (*Sox*, *Fox*, *Pax*), immuno-genes (NACHT, SRCR, TLR families), and nervous-system families (*rhol*, opsin), were all markedly expanded in those species, but not in *M. globulus*.

To place these copy number patterns in functional context, we examined the expression of the same curated gene families across adult *M. globulus* tissue transcriptomes (**Figure 5C**). Several families showed clear preferential expression in particular tissues. Consistent with their known function, biomineralisation families (MSP130, SM30, SM50) were relatively enriched in spine and tube feet, whilst showing relatively lower expression in the ovary and gut. Pigmentation-associated families (PKS, FMO) were relatively enriched in the spine and gut, and reduced in the ovary. Transcription factors and nervous system associated families (SOX, fox, pax, rhol, opsin) were generally highest in the spines, with several elevated in the gut, while the ovary was consistently depleted. Since the heatmap was row-normalised, this data indicates relative tissue bias within each gene family rather than absolute expression differences among families.

The genome-wide CAFE (v5) analysis revealed that gene family turnover showed variation across the echinoid tree (**Figure 5A**). *M. globulus* in particular exhibited the fewest expansions of the species studied and a comparatively large number of contractions. When examining the distribution of gene family copy numbers, *M. globulus* stood out once more for its high proportion of single-copy genes, and lower proportion of multi-copy and duplicate genes (**Figure 5B**). These numbers suggest that *M. globulus* diverged before the extensive gene duplication events shared by several camarodont urchins belonging to the Infraorder Echinidea (Marlétaz et al., 2023) and may possess a less reshuffled gene repertoire.

In line with our previous analyses, *M. globulus*, once again exhibited a conservatively evolving gene repertoire, with contractions across most of the targeted functional categories. Notably, SRCR and TLR families, both components of the echinoid immune system (Barela Hudgell and Smith, 2021; Pancer, 2001), show relatively reduced copy numbers in *M. globulus* compared to other echinoids, indicating a possible reduction in immune gene number. SRCR genes are massively expanded in the holothurian *H. leucospilota*, suggesting that the large SRCR repertoires arose early in echinozoan evolution, with subsequent lineage-specific expansions and contractions. This pattern mirrors the possibility raised that the reduced gene complement in *M. globulus* may reflect retention of a more ancestral state rather than secondary contraction.

Overall, the gene family landscape of *M. globulus* points to a streamlined genomic architecture, in contrast to the dynamic gene family expansions in the camarodonts belonging to the Infraorder Echinidae. The absence of expansion across all functional categories suggests that *Mespilia*’s distinctive morphology and pigmentation has not evolved through duplication-driven novelty. The relative lack of gene expansion in *M. globulus* could represent an important feature for subsequent gene functional analysis. When using Cas9 or MASOs for gene knockout/knockdown, having single or minimal family members enhances knockout/knockdown strategies and simplifies functional interpretations.

## Discussion

Recent advances in genome editing and functional genomics have increased the need for model organisms with well-annotated genomes and accessible life cycles well adapted to aquarium environments. Here, we present the tuxedo urchin *M. globulus* as a promising echinoid model. It offers rapid development, easy aquarium adaptability, and multiple chromosome-scale genome assemblies. These traits enhance established sea urchin models like *S. purpuratus, P. lividus,* and *L. pictus*, expanding the tools available for echinoderm research.

One major benefit of *M. globulus* is its quick and consistent life cycle under controlled aquarium conditions. Embryonic development is fast, with larvae reaching metamorphosis in about two weeks and achieving sexual maturity in four to six months. This timeline is much shorter than that of several common temperate sea urchin models and is similar to *Lytechinus.* However, the higher degrees of gene duplication in *Lytechinus* make it potentially less-amenable to functional study than *M. globulus,* which our gene family analyses indicate has a comparatively streamlined gene set. Across various functional categories, this species shows fewer gene-family expansions and a higher proportion of single-copy genes than a number of other sequenced sea urchins. Instead of indicating significant gene loss, this pattern may suggest the retention of a more ancestral gene complement. This reduced redundancy in gene families might help in interpreting functional perturbation experiments, as gene disruptions are less likely to be compensated by related copies.

Further, our results also show that *M. globulus* works well with experimental methods commonly used in developmental and functional genomics. Hybridization chain reaction effectively detected spatial expression of key regulatory genes, and CRISPR-Cas9 genome editing produced predictable phenotypic outcomes when targeting polyketide synthase. Together, these findings suggest that molecular and genetic methods used in other echinoid models can be easily applied to this species, establishing it as a tractable model for developmental biology research.

Chromosome-scale assemblies of the red and blue morphs further demonstrate the value of *M. globulus* as an experimental system. Both assemblies recover the usual set of 21 echinoid chromosomes and show strong conservation of gene order. Whole-genome comparisons revealed high sequence similarity between morphs and little evidence of large-scale structural rearrangements. Any observed differences in chromosome orientation result from the arbitrary arrangement of assembled scaffolds rather than structural variation. Comparative analyses also position the *M. globulus* genome within the larger context of echinoid evolution. Echinoderm genomes do show lineage-specific rearrangements (Marlétaz et al., 2023), but the tuxedo urchin maintains a relatively stable chromosomal structure and shares ancestral fusion events typical of euechinoids. The lack of divergence in chromosome-wide coverage depth does not support the presence of a differentiated sex chromosome. Rather, it indicates a largely shared genome between sexes, with localized deviations from sex parity. This indicates that sex determination in sea urchins is not necessarily controlled by a single sex chromosome, but is likely polygenic. This would also be consistent with a system in which sex chromosomes are evolving, but are not yet differentiated (Graves, 2015).

Lastly, the widespread cultivation of *M. globulus* in the marine aquarium trade offers a practical advantage for experimental research. These urchins are often produced through aquaculture instead of wild collection. This ensures consistent availability while reducing pressure on natural populations. Captive breeding may also lower genetic variability compared to wild-caught individuals, possibly enhancing experimental reproducibility.

These features establish *M. globulus* as a practical echinoid model for functional and comparative genomics. Its fast life cycle, compatibility with modern molecular tools, and relatively streamlined genome create a valuable platform for exploring the genetic and developmental foundations of echinoderm biology.

## Materials and methods

### Adult Culture system

The blue *Mespilia globulus* specimen was sourced from the culture maintained at the Horniman museum and was originally provided by the Tropical Marine Centre (TMC) which imported them from Indonesia. The red sample was sourced from Ocean Reefs and Aquaria, (orafarm.com). Urchins were acclimated to controlled conditions in a quarantine tank for one week before introduction to the main tank system. The tank system consisted of three 100 L tanks with a temperature of 24°C and salinity of 35 parts per thousand (ppt). Reverse osmosis (RO) water is automatically added to compensate for evaporation and a protein skimmer removes dissolved organic waste.

Adult urchins were fed twice a week with a mixture of rehydrated kombu (or Ulva) and carrots. Once a week, tanks are cleaned and pH (8.0-8.4), Nitrate (0-20ppm), Nitrite (0-0.5ppm) and ammonia levels (0-0.25ppm) monitored. The system was maintained in a 8:16-hour light-dark cycle with LED lighting.

### Spawning and larval rearing

Spawning was induced using heat shock, or by injection of 100 µL of 0.5M KCl or 2mM acetylcholine in filtered sea water into the coelom of the adult. For heat shock induced spawning, adults were brought to 29 °C for up to 45 minutes and then transferred back to 24 °C. If unsuccessful, spawning is induced by injection with KCl (Nesbit et al., 2019). Eggs were fertilized by adding 100 µL of freshly diluted (1,000x) sperm, and then filtered through a 125 µm mesh and maintained in 5 L of sterile filtered (Sartopore, 0.2 µm) artificial (H2Ocean) seawater (ASW) at a density of 1 egg per 4 mL with aeration. From the pluteus stage, 50 µL of *Rhodomonas salina* culture (Scottish Association for Marine Science, SAMS) was added to the culture twice a day, and motor-mounted paddles were inserted into the culture to maintain a smooth agitation to keep larva in suspension as described by (Nesbit et al., 2019) (Figure 2).

Approximately two weeks post-fertilization, larvae underwent metamorphosis and settled on the beaker floor. They were then carefully transferred using a fine paint brush into a large, aerated plastic tub (30×50×30 cm) and the juveniles were fed the diatom *Nitzschia ovalis* daily until they became visibly distinguishable by their distinct reddish-purple pigmentation. After acclimation, they were maintained in culturing vessels in the main broodstock tank where their diet is expanded to include a combination of *N. ovalis* and kombu until they reached sexual maturity, which takes approximately 4 months post-metamorphosis (mpm), at which time they could be transferred to the main tank.

### Algal culture

Base f/2 algae growth media (Florida Aquafarms Inc.) is prepared in autoclaved artificial seawater (ASW) at 30ppt NaCl. *R. salina* cultivation flasks are sealed and continuously aerated using a serological pipette attached to an air pump (Figure 2). *N. ovalis* cultivation flasks are sealed and placed on a shaker at 35 rpm. The cultures are maintained under a 12h/12h light/dark cycle at 22°C and fertilised with 0.5ml of f/2 twice a week. When cultures reach a dense growth state, they are diluted at 1:3 with base media to maintain optimal growth conditions.

Algal cultures are harvested by collecting 50 mL aliquots, centrifuged and washed 3 times in filtered artificial seawater (FASW). Culture density was estimated by cell counting and used to determine the required volume of algae for larval rearing, as detailed in Supplemental Table 1.

### Fluorescent & HCR staining

Specimens were fixed in 4% PFA for 2 hours at room temperature (RT), then washed 3x 5min in FASW. Samples were stained overnight with Phalloidin (Invitrogen) 25µL in 1 mL of FASW and mounted in EVERBRIGHT (Biotium) medium with DAPI. Slide edges were sealed with nail polish to preserve the preparation.

For Hybridization Chain Reaction (HCR) fluorescent *in situ*, samples were fixed in 4% PFA for 30 minutes at RT, washed, and dehydrated in graded methanol series before storage in 100% methanol. Prehybridization was conducted at 37°C for 3min followed by overnight hybridization also at 37° with 8 nM probe concentration. Amplification for hairpin activation involved denaturation at 95°C for 90 seconds, cooling at RT and overnight amplification at 25°C with a 40 nM hairpin concentration. Samples were washed in SSC buffer and PBS Tween-20. Specimens are counterstained with DAPI (1 µg/mL) and mounted in glycerol before confocal imaging.

HCR probes were designed using the HCR Probe Maker CL (*HCRProbeMakerCL*, n.d.): https://github.com/rwnull/HCRProbeMakerCL) and were then ordered from Integrated DNA technologies (IDT). Amplifiers and other HCR solutions were purchased from Molecular Instruments.

### Crispr/Cas9 in Tuxedo urchin

Three gRNAs were designed against Mg PKS1 using CRISPRscan software (https://www.crisprscan.org/) and ordered from IDT (https://www.idtdna.com/page): MgPKSsgRNA1 (CTGCAAAGGAGGGACCCTTG), MgPKSsgRNA2 (ACTTAACGGTTTGAGGACTA) and MgPKSsgRNA3 (CGGGGACCCTCTTGAGGCAG). They were received as powder and were resuspended in nuclease free water to 100 uM. The Cas9 protein was obtained from IDT (Alt-R™ S.p. Cas9 Nuclease V3, 500 µg, 1081059). The stocks of gRNAs and the Cas9 protein can be stored long term at −20 degrees. The injection mixes were produced as follow: 3.2 ul of nuclease free water, 2.5 ul of gRNA (for 3 gRNAs: 0.83 ul of each gRNA), 0.3 ul of Cas9 protein IDT (stock at 62uM from company so we get about 2uM final). These mixes were centrifuged at 4000 rpm for 30 seconds at room temperature and incubated for 5 minutes at room temperature to pre-bind the Cas9 and the gRNAs together. The FITC conjugated dextran dye (Fisher Scientific, D-1821) was then added as an injection reporter, the stock tube is made at 5 mM, filtered using the centrifugal filter units (Millipore, UFC30GV0S) and centrifuged for 1 minute at 13000 rpm at room temperature, and 2 ul of this dye was added to the injection mix. The final mixes were centrifuged for 1 minute at 13 000 rpm at room temperature and kept on ice. The control mix contains the Cas9 protein and no gRNA.

The *M. globulus* eggs were kept at room temperature, and dejellied by filtering them 5 times through a 100 uM nylon mesh (Fisherbrand, 22363549). They were aligned on a protamine sulfate (1% solution) pre-coated plate containing sea water with PABA 2 mM to prevent the hardening of the fertilization envelope during the injection. PABA stands for 4-aminobenzoic acid (Sigma, A9878), a 20 mM stock is made in sea water at pH 8, and can be kept at 4 degrees for several weeks. Eggs were fertilized with diluted sperm (the stock of concentrated sperm was kept at room temperature and diluted by 200 times in sea water before use). Eggs were injected after fertilization.

For the injection, glass capillaries with filament were purchased (Narishige, GD-1), and pulled with the needle puller (Narishige model PC-10). 0.5 ul of injection mix was loaded into a needle. The parameters for microinjection were controlled with the Femtojet 4i (Eppendorf). After injection, the fertilized eggs were rinsed two times with sea water without PABA. The embryos were cultured at room temperature and imaged at larva stage using the AxioPlan2 microscope connected to the Axiocam 820 color camera.

### Genome Assembly

DNA was extracted from flash-frozen gonadal tissue of a single *M. globulus* individual using the Nanobind tissue kit (Pacific Bioscience). DNA molecular weight was verified using a Femto Pulse instrument before HiFi library preparation. Briefly, the DNA was sheared to obtain a narrow size profile (15-18 kb) before sequencing adapter ligation and circularization. Multiple sequencing rounds were performed, with a consensus computed using the Circular Consensus Sequencing (CCS) tool from PacBio. Genome assembly was conducted using Hifiasm with default parameters. Haplotypes were then filtered using the ‘purge_dups’ tool (https://github.com/dfguan/purge_dups) yielding a haploid assembly.

A Hi-C library was constructed from M. globulus gastrula (24 hpf) using the Arima-HiC kit. Briefly, frozen tissue was pulverised in liquid nitrogen and chromatin crosslinked with PFA before restriction enzyme digestion. Following biotinylation and proximity ligation, DNA was fragmented and biotinylated fragments were enriched for Hi-C sequencing library construction.

Hi-C sequencing reads were mapped to the reference genome using bwa-mem (v0.7.17). Ligation junctions were identified, and PCR duplicates were removed using Pairtools (v1.02). The processed reads were then used to generate a .bam file, which was used as input to generate a Hi-C contact map in .hic format using Juicer (v1.6). This contact map facilitated scaffolding of the genome assembly. For scaffolding, we tested YAHS (v1.2). YAHS provided a more contiguous assembly, which was subsequently curated and adjusted manually using Juicebox before a final assembly was generated (**Supp. Table 1)**.

### Genome annotation

The genomes of *M. globulus* were annotated using a previously described pipeline (https://github.com/eparey/AnnotateSnakeMake) (Parey et al., 2024) incorporating transcriptomic data from 24 newly generated samples (Supplementary Table 1).

Gene Ontology (GO) annotations for *M. globulus* genes were generated using eggnog-mapper (v2.1.12). Functional enrichment analysis was performed using the enricher function from the ClusterProfiler R package (Yu, 2018), employing hypergeometric tests to identify significantly enriched GO terms. The resulting enrichment data was summarized using REVIGO, and top ontology terms were selected based on their ‘dispensability’ scores to identify the most representative and non-redundant functional categories.

### Sex-biased region

DNA from a male and a female tuxedo sea urchin from the blue morphotype were extracted using the Nanobind PanDNA kit (Pacbio), paired-end library were built using Truseq kit (Illumina) and sequenced on an Illumina Novaseq X at 30x coverage. Reads were aligned to the blue male and red female assemblies using bwa-mem2 (v2.2.1) and coverage depth was calculated in 10kb windows using Mosdepth (v0.3.14) (Pedersen and Quinlan, 2018). SNP calling was carried out using bcftools (v 1.23.1) with a minimal mapQ of 20. Normalisation, statistical analysis and plotting were performed in R.

### Synteny analyses

Pairwise orthologues were derived from Orthofinder between species reported in Table S5 and their position along chromosomes indexed. Reciprocal locations of orthologues were compared and plotted with custom code and statistically assessed using Fisher test as previously described (Parey et al., 2024). Synteny linear plots as in Figure 3C were assembled using the Rideogram package in R.

### RNA Extraction

Extraction of RNA from embryos and larvae was performed using the RNeasy Plus Mini Kit (Qiagen), following the manufacturer’s recommended protocol. Embryos, larvae, and juveniles were homogenised in RLT plus buffer after addition of beta-mercaptoethanol using a Cryolyser (Stretton Scientific). gDNA was removed using the gDNA eliminator spin column. RNA integrity was verified using the Tapestation instrument (Agilent). RNA-seq libraries were prepared using the TruSeq kit (Illumina) and sequenced with 30M paired-end reads. Extraction of RNA from adult tissues (ovary, spines, tube feet, and gut) was accomplished by homogenization in Trizol and purification of the RNA following the manufacturers protocol.

### Gene family expansion/contraction analysis

Gene family turnover across the echinoid phylogeny was modelled using CAFE (v5)(Mendes et al., 2021), which estimates rates of gene family gain and loss under a birth-death model. Gene families were inferred with OrthoFinder (v3.0.1)(Emms and Kelly, 2019), which was run on the following species with default parameters: *Holothuria leucospilota*, *Mespilia globulus*, *Paracentrotus lividus*, *Strongylocentrotus purpuratus*, *Lytechinus variegatus*, and *Lytechinus pictus* (**Supplementary Table 1**).

Protein phylogeny was inferred from 1,642 single-copy 1:1 orthogroups. Sequences were aligned with MAFFT (v.7.520) and filtered to remove poorly aligned sites. The maximum-likelihood tree was inferred with RAxML-NG (v1.2.2) under the LG+F+G4 substitution model. Tree searches were performed using 10 starting trees, and branch support was assessed with 100 bootstrap replicates. Bayesian phylogenetic analyses and divergence-time estimation were conducted using PhyloBayes (v4.1) under the CAT+GTR model. Chains were run until > 5,000 iterations, inspected for convergence, and summarised with 500 burn-in.

The OrthoFinder gene count matrix and the time-calibrated ultrametric tree were input into CAFE (v.5.0), a tool which estimates the global birth-death parameter (λ) by maximum likelihood from the data. Orthogroups with extremely high copy number variation (>100 copies in any species) were analysed separately to prevent bias in rate estimation, using the λ value derived from the filtered dataset. Gene families with family-wide p<0.01 were considered significantly evolving and classified as expanded or contracted. The number and proportion of expansions, contractions, and no change, were summarised per species and visualised on the species tree (**Figure 5A**).

To assess the evolution of specific gene families associated with key echinoid traits, we assembled a curated set of functionally characterised genes representing pigmentation, biomineralisation, immune function, transcriptional regulation, and neural development. Multiple representative protein sequences from *Strongylocentrotus purpuratus* and other echinoids were collected for each functional group from Echinobase and NCBI and were matched to their corresponding orthogroups in the OrthoFinder output.

Gene copy numbers for each relevant orthogroup were extracted for each species and normalised by z-scores to visualize relative expansions/contractions. The functional categories included pigmentation (PKS, FMO), biomineralization (SM30, SM50, MSP130), immune system (SRCR, NACHT, TLR), transcription factors (SOX, fox, pax), and nervous system function (opsin, rhol). Heatmaps were generated to compare copy number variation across species, and orthogroups identified as significantly evolving in CAFE were annotated accordingly.

To relate gene family evolution to tissue-level expression in *Mespilia globulus*, we examined the same curated set of multifunctional gene families in adult tissue transcriptomes from tube feet, spine, ovary, and the gut. Expression values were transformed as log2(TPM+1) and row-normalised as z-scores to visualise the relative tissue enrichment in each family. All heatmaps were generated in Python using the seaborn package (Waskom, 2021).

### SEM

Tests were isolated from spines, tube feet, and pedicellaria, rinsed in deionized water, and airdried before sputter coating with a gold/palladium 60:40 mix and imaged with a Thermo Apreo VS Scanning Electron Microscope.

## Data availability

The genome of red and blue tuxedo genomes as well as the corresponding sequencing data have been deposited to NCBI under the Bioproject accession PRJNA1477966. Code and datasets used for analysis have been deposited in this repository: https://github.com/fmarletaz/tuxedo

## Author contributions

F.M., O.M., J.R.T., J.C. and G.W. initiated the work on *M. globulus* as a model. F.M., O.M. and G.W. conceived this project. O.M. and M.E.M. established culture protocols. N.O. and G.W. performed CRISPR experiments, collected tissues and specimens. L.K. performed gene family and transcriptomic analyses. C.S. and G.B. took care of urchin husbandry. G.B. and L.P. performed HCR staining. E.P., O.M. and F.M. assembled and annotated the genomes.O.M. and F.M. performed sex comparison and synteny analyses. O.M., F.M. and G.W. wrote the manuscript with input from all authors.

## Acknowledgments

This work was supported by a Royal Society URF fellowship to F.M. supporting O.M. (URF\R1\191161 and URF\R\241014). L.K., L.P., C.S. and M.E.M. were supported by a BBSRC research grant (BB/V01109X/1) and a Leverhulme grant (RPG-2021-436) to F.M. E.P. was supported by a Royal Society Newton Fellowship (NIF\R1\222125), and a Leverhulme Research grant (RPG-2025-274). J.R.T. was supported by a Leverhulme Trust Early Career Fellowship. G.W. and N. O. were supported by the United States National Institutes of Health (1R35GM140897, G.M.W.)

